# Silencing of transposable elements may not be a major driver of regulatory evolution in primate induced pluripotent stem cells

**DOI:** 10.1101/142455

**Authors:** Michelle C. Ward, Siming Zhao, Kaixuan Luo, Bryan J. Pavlovic, Mohammad M. Karimi, Matthew Stephens, Yoav Gilad

**Author notes:** Correspondence should be addressed to M.C.W, and Y.G.

## Abstract

Transposable elements (TEs) comprise a substantial proportion of primate genomes. The regulatory potential of TEs can result in deleterious effects, especially during development. It has been suggested that, in pluripotent stem cells, TEs are targeted for silencing by KRAB-ZNF proteins, which recruit the TRIM28-SETDB1 complex, to deposit the repressive histone modification H3K9me3. TEs, in turn, can acquire mutations that allow them to evade detection by the host, and hence KRAB-ZNF proteins need to rapidly evolve to counteract them. To investigate the short-term evolution of TE silencing, we profiled the genome-wide distribution of H3K9me3 in induced pluripotent stem cells from ten human and seven chimpanzee individuals. We performed chromatin immunoprecipitation followed by high-throughput sequencing (ChIP-seq) for H3K9me3, as well as total RNA sequencing. We focused specifically on cross-species H3K9me3 ChIP-seq data that mapped to four million orthologous TEs. We found that, depending on the TE class, 10-60% of elements are marked by H3K9me3, with SVA, LTR and LINE elements marked most frequently. We found little evidence of inter-species differences in TE silencing, with as many as 80% of orthologous, putatively silenced, TEs marked at similar levels in humans and chimpanzees. Our data suggest limited species-specificity of TE silencing across six million years of primate evolution. Interestingly, the minority of TEs enriched for H3K9me3 in one species are not more likely to be associated with gene expression divergence of nearby orthologous genes. We conclude that orthologous TEs may not play a major role in driving gene regulatory divergence between humans and chimpanzees.

## Introduction

Over half of primate genomes are annotated as transposable elements (TEs) (Jurka 2000; de Koning et al. 2011). Primate TE sequences reflect their evolutionary history; some TEs are conserved over multiple phylogenetic clades and orders, whereas others are restricted to particular lineages. Considering the degree of sequence similarity between the human and chimpanzee genomes, it is not surprising that many of the same TEs are present in both species (Chimpanzee and Analysis 2005; Ramsay et al. 2017). In general, TE activity, especially from endogenous retroviruses (ERVs), has declined in the hominid lineage relative to other mammalian lineages (Lander et al. 2001). However, a fraction of evolutionarily recent TEs are active in the human genome, including HERV-K ERVs (Tonjes et al. 1996; Medstrand and Mager 1998; Fuchs et al. 2013), members of the Alu (Batzer and Deininger 1991; Batzer et al. 1991), L1 (Kazazian et al. 1988; Brouha et al. 2003), and SVA (Ostertag et al. 2003; Wang et al. 2005) families. Most studies on mammalian TEs have focused on humans and mice, with a handful that describe TE activity in the ERV and L1 families in chimpanzees (Yohn et al. 2005; Marchetto et al. 2013; Mun et al. 2014).

Because of their ability to newly insert into the genome, TEs are considered a potential source of regulatory innovation (Kazazian 2004; Cordaux and Batzer 2009). Indeed, TEs can serve as primate-specific regulatory sequence (Jacques et al. 2013), act as enhancer elements (Lynch et al. 2011; Chuong et al. 2013; Xie et al. 2013), carry transcription factor binding sites (Wang et al. 2007; Bourque et al. 2008; Kunarso et al. 2010; Schmidt et al. 2012), and contribute themselves to the transcriptome (Faulkner et al. 2009; Kelley and Rinn 2012; Kapusta et al. 2013). The influence of TEs on transcriptional programs has been studied in many broad contexts from the immune system (Chuong et al. 2016) to pregnancy (Lynch et al. 2015), to pluripotency (Macfarlan et al. 2012). Yet, ultimately, the regulatory impact of TEs has been studied through anecdotal examples. In particular, there are no published comparative genome-wide surveys of the regulatory effects of TEs in the context of primate evolution.

Due to the potential deleterious effects of new TE insertions, especially in the context of developmental regulatory programs, TE activity is thought to be tightly controlled by host repressive machinery. In particular, it was suggested that in embryonic stem cells (ESCs), KRAB-containing Zinc Finger genes (KRAB-ZNFs) can recognize TEs in a sequence-specific manner, recruit the co-factor TRIM28 (also known as KAP1), and histone methyltransferase SETDB1 (also known as ESET), to effect silencing through the deposition of the repressive histone three lysine nine trimethylation (H3K9me3) modification (Schultz et al. 2002; Wolf and Goff 2009; Matsui et al. 2010; Rowe et al. 2010; Friedli et al. 2014; Turelli et al. 2014; Najafabadi et al. 2015). After the establishment of TE silencing by H3K9me3, TEs are subjected to de novo DNA methylation, which persists in somatic and germ cells (Bourc'his and Bestor 2004; Karimi et al. 2011; Rowe and Trono 2011; Quenneville et al. 2012; Rowe et al. 2013; Smith et al. 2014; Barau et al. 2016; Walter et al. 2016). Some TEs can evade detection by host KRAB-ZNFs by acquiring mutations. It was thus hypothesized that the host repressor mechanisms need to evolve rapidly in order to maintain genome integrity (Huntley et al. 2006; Thomas and Schneider 2011; Kapopoulou et al. 2016).

TEs are silenced by KRAB-ZNFs particularly in early embryonic development (Chuong et al. 2017); however it is difficult to obtain biological material from humans and other primates to determine how this process evolves. Induced pluripotent stem cells (iPSCs) provide a useful, ethical model system of the embryonic inner cell mass to study this phenomenon. This cell type is of further interest as we know of examples of primate-specific TEs that are not silenced, but actively transcribed and functionally relevant in this cell type in human (notably HERVH elements) (Macfarlan et al. 2012; Santoni et al. 2012; Fort et al. 2014; Lu et al. 2014; Wang et al. 2014; Grow et al. 2015). This might imply specificity of TE regulatory mechanisms, and hints at the fact that evolutionarily recent TE sequence could play a role in differentiating gene regulatory networks between species.

The establishment of a panel of non-retroviral reprogrammed iPSCs from multiple humans and chimpanzees has allowed us to comprehensively characterize the extent of TE silencing in both species (Gallego Romero et al. 2015; Banovich et al. 2016; Burrows et al. 2016). In order to gain insight into how TE regulation, specifically TE silencing, evolves between humans and chimpanzees, we globally profiled the predominant output of targeted silencing mechanisms in pluripotent cells: the histone modification H3K9me3. Our goals were to identify inter-species differences in TE silencing patterns, characterize gene regulatory divergence that can potentially be explained by differences in TE silencing between species, and develop hypotheses regarding the mechanisms that underlie such inter-species differences in TE regulation. As we report in detail, our observations led us to conclude that TE silencing is remarkably conserved in humans and chimpanzees, and that – at least in our comparative iPSC system, TEs do not play a major role as drivers of gene regulatory divergence between these two closely related species.

## Results

To perform a comparative study of TE silencing, we characterized genome-wide distributions of the repressive histone modification H3K9me3 in iPSCs from seven chimpanzees and ten humans (Fig. 1A, Table S1). The iPSCs lines we used were deeply characterized in this study and previously (see Methods; and (Gallego Romero et al. 2015; Banovich et al. 2016; Burrows et al. 2016)). Briefly, all cell lines were confirmed to have a normal karyotype (Fig. S1A), have the ability to form embryoid bodies that consist of cells from the three main germ layers (Fig. S1B), display global gene expression signatures consistent with pluripotent cells (Fig. S2), and express key pluripotency factors detectable by immunocytochemistry (Fig. S3). All samples were processed in species-balanced batches at each experimental step (Table S2).

**Figure 1:**
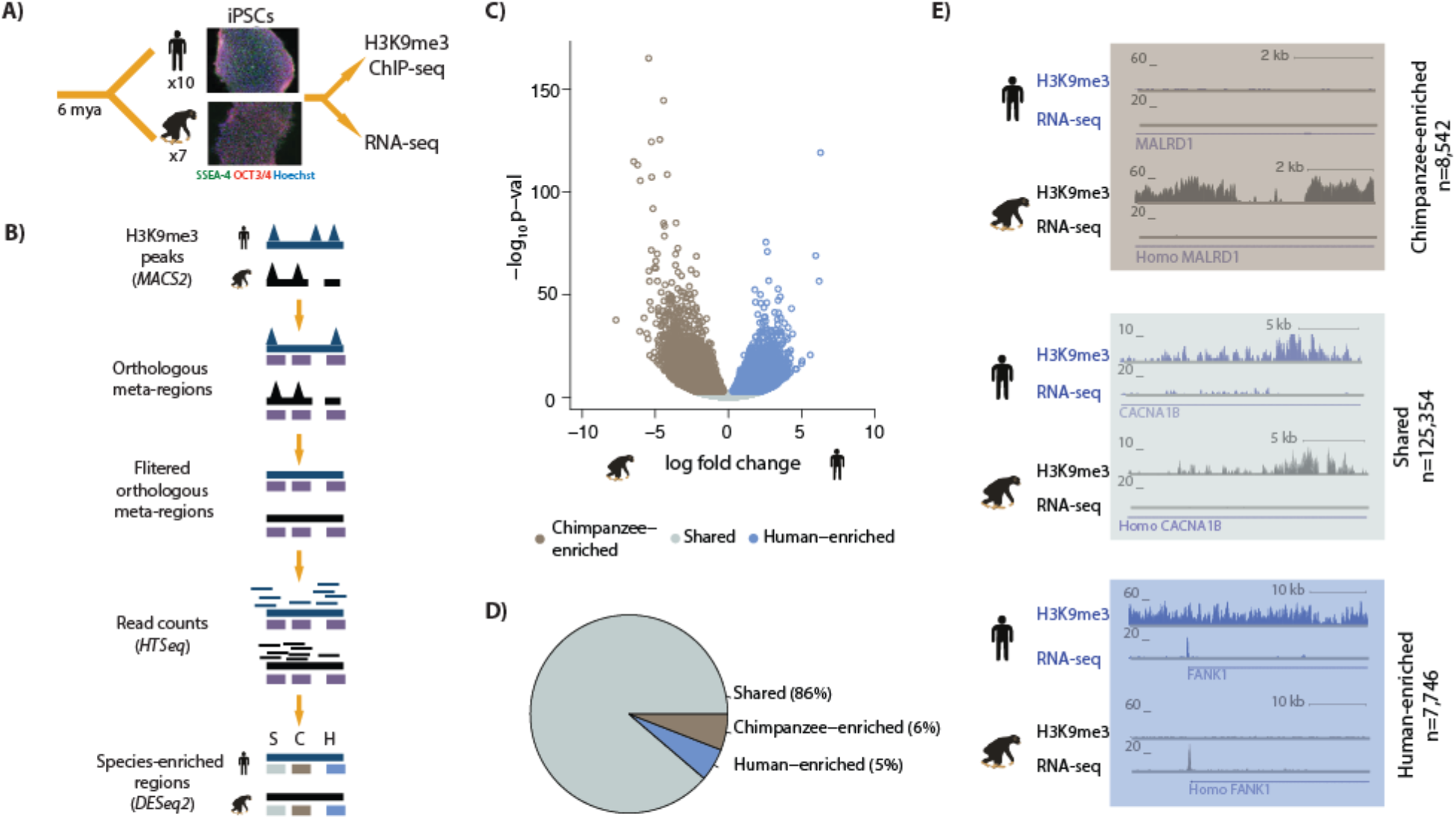
Majority of H3K9me3 regions are similarly enriched in human and chimpanzee iPSCs. (A) Experimental design of the study. (B) A heuristic description of the computational pipeline to identify regions of differential H3K9me3 enrichment between humans and chimpanzees. Human genome, ChIP-seq peaks and ChIP-seq reads (blue); chimpanzee genome, ChIP-seq peaks and ChIP-seq reads (black), orthologous H3K9me3 regions (purple). (C) Volcano plot representing the regions similarly enriched in both species (grey: Shared), enriched in human (blue), or enriched in chimpanzee (brown) at 1% FDR. (D) Pie-chart representing the proportion of regions within the Shared, Human-enriched and Chimpanzee-enriched categories. (E) Examples of individual loci representing each of the three defined categories. The top track represents sequencing reads from H3K9me3 ChIP-seq experiments, and the bottom track gene expression levels of the *MALRD1, CACNA1B* and *FANK1* genes determined by RNA-seq.

We characterized H3K9me3 in the 17 iPSC lines by performing chromatin immunoprecipitation followed by high-throughput sequencing (ChIP-seq). In order to increase genome-mapping efficiency, paired-end sequencing libraries were prepared, and only high quality, properly paired reads retained (see Methods and Fig. S4 and Table S3). As a first measure of quality of our data, consistent with previous reports, we observed an enrichment of H3K9me3 ChIP-seq reads at ZNF genes in both species (Fig. S5)(Roadmap Epigenomics et al. 2015).

As a first step of our analysis we identified regions of broad H3K9me3 ChIP enrichment in each individual independently (at an FDR of 10%; see Methods and Fig. S6-7). In order to compare H3K9me3 enrichment across species, we focused exclusively on reciprocally best match orthologous regions in human and chimpanzee (see Methods). We excluded from our data peaks that mapped outside those orthologous genomic regions (approximately 10% of the ChIP-seq data were excluded; Fig. S8 and Table S3). We further filtered the data based on overall genome mappability analysis (using a cutoff of 0.8 in both species; see Methods), to exclude regions in which sequencing reads can be mapped to one species at a substantially higher probability than the other. We then defined orthologous H3K9me3 ChIP-seq regions as those where a ChIP-seq peak, contained within orthologous human-chimpanzee genomic regions, was identified in at least one individual, in either species in our study (see Methods). We refer to these as orthologous ChIP-seq regions (rather than ChIP-seq peaks), because these regions are defined across all individuals, while the ChIP-seq peak may have been identified in only a subset of individuals.

The definition of ChIP-seq regions facilitates a quantitative comparison of H3K9me3 enrichment across species. To do so, we no longer relied on ChIP-seq peaks (which are classified with incomplete power in each individual), but rather we calculated the number of sequencing reads that were mapped to each orthologous region in each individual. In other words, once the orthologous regions were defined based on the classification of a ChIP-seq peak in at least one individual (using an arbitrary statistical cutoff), we considered counts of reads mapped to these regions from all individuals in our subsequent comparative analysis. By using this approach we sidestep the difficult challenge of accounting for incomplete power to detect ChIP-seq peaks in each individual or species independently. To avoid considering regions where we clearly have insufficient power to compare read counts across species, we performed a minimal pre-filtering step and only included the 150,390 regions with at least one read count in more than half of the individuals (regardless of species).

We considered overall properties of the read count data in the ChIP-seq orthologous regions. As expected, data from different individuals cluster by species (with the exception of one human individual, which therefore was removed from subsequent analysis; Fig. S9-10), and inter-species variation in read counts is greater than intra-species variation (Fig. S11). We then used the framework of a linear model with a fixed effect for species to identify orthologous ChIP-seq regions with differences in H3K9me3 enrichment between humans and chimpanzees (Fig. 1B, Supplemental Table 1, and Methods). At FDR < 0.01, we found 16,288 (11% of the tested 141,642 regions) differentially enriched regions between the two species (Fig. 1C-D, Fig. S12, and see examples in Fig. 1E). The proportion of differentially enriched regions is robust with respect to our data filtering steps (Fig. S13), and is consistent across two independent methods (Table S4A). The majority of regions (81 or 87% depending on the method used) have an effect size (log fold change between species) of less than one (Fig. S14 and Table S4B).

### Orthologous TEs tend to be silenced more often than species-specific TEs

To compare TE silencing by H3K9me3 across species, we identified a comprehensive set of 4,248,188 orthologous TEs in the human and chimpanzee genomes (see Methods and Fig. 2A, Fig. S15). We considered the ChIP-seq data in the context of the orthologous TEs, and focused on the 12% of orthologous TEs that overlap an orthologous H3K9me3 region with at least 50% of their length (Fig. 2B and Supplemental Table 2; we note that our subsequent conclusions are robust with respect to the specific cutoff used to classify a TE as overlapping with H3K9me3 regions). Consistent with previous reports (Karimi et al. 2011; Turelli et al. 2014), we found enrichment of LTRs (Pearson’s Chi-squared test; *P* < 0.001), and SVA elements (*P* < 0.001), among TEs that overlap H3K9me3 regions (Fig. 2C).

**Figure 2:**
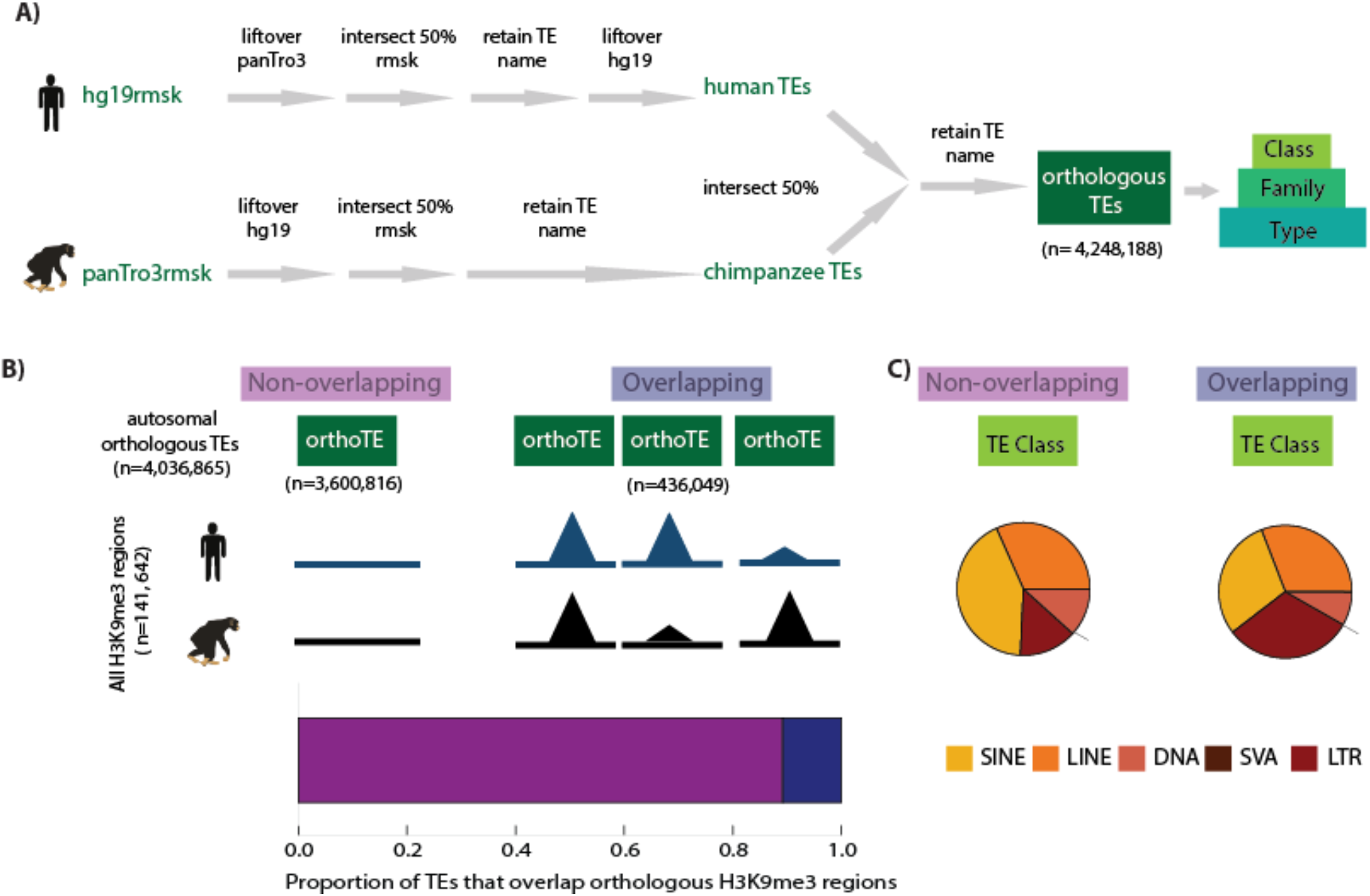
Transposable element silencing is associated with particular transposable element classes. (A) Bioinformatic pipeline used to identify TEs that are orthologous between humans and chimpanzees using RepeatMasker (rmsk) tracks in each species. Each TE instance is hierarchically classified into Class, Family and Type. (B) The proportion of all TEs that overlap an orthologous H3K9me3 region (H3K9me3 peak identified in at least one individual of one species) (dark blue) compared to those that do not overlap (magenta). (C) The proportion of all orthologous TEs belonging to the five main TE classes (SINE: yellow, LINE: orange, DNA: peach, SVA: brown, LTR: red) that do not overlap an H3K9me3 region in either species, and the proportion of TEs that do overlap a H3K9me3 region. The small number of SVA elements means that this class is not visible in the pie chart.

Given the proposed evolutionary arms race between TEs and the host repressive machinery (Jacobs et al. 2014), we sought to investigate the within-species patterns of silencing at both orthologous and non-orthologous TEs. We further considered TEs whose type exists in only one of the species (for example, SVA-E & SVA-F, which are human-specific TE types; Table S5). Generally, we found that the same TE classes that most frequently overlap with H3K9me3 regions in the orthologous TE set, also most frequently overlap with H3K9me3 in the non-orthologous TE set (see Methods; Fig. 3). Yet, there are specific TE classes within each species that show a clear reduction in the proportion of TEs overlapping H3K9me3 regions in the non-orthologous, and species-specific TE types (Pearson’s Chi-squared test for both species; *P* (SVA) < 0.001; *P* (LINE) < 0.001; Fig. S16). We also found reduction in overlap in the LTR class in human (*P* < 0.001), however a significant increase in overlap is observed in chimpanzee-specific LTRs (*P* < 0.001; Fig. 3). Notably there are many more species-specific LTRs in chimpanzee (10,798), than in human (534). This result could intuitively be interpreted as suggesting that more recent TEs are generally less likely to be silenced, but our subsequent analysis may indicate otherwise.

**Figure 3:**
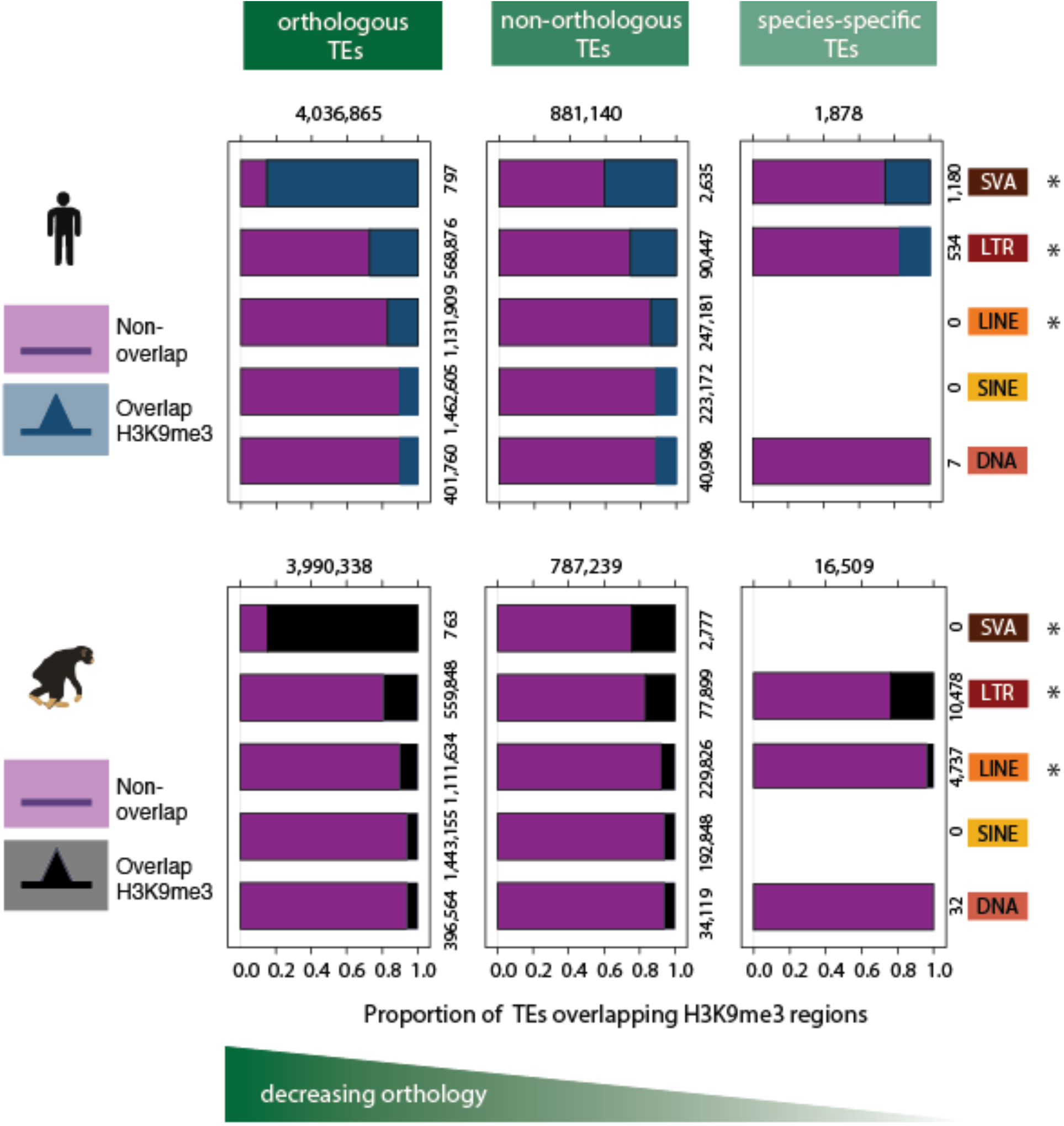
Patterns of preferential TE class-based silencing are maintained as orthology decreases. Within-species analysis of the proportion of orthologous TEs (left panel), TEs that are not orthologous between the two species (middle panel), and TEs that are not orthologous and whose type is annotated in a single species (right panel), that overlap any H3K9me3-enriched region identified in that species. Magenta represents those TEs that do not overlap with any H3K9me3 region identified in that species, blue represents TEs that overlap any H3K9me3 regions in human, and black those TEs that overlap with H3K9me3 regions in chimpanzee. Asterisk denotes when there is a significant difference between orthology categories (Pearson’s Chi-squared test; *P* < 0.001).

### Majority of orthologous TEs are similarly silenced in human and chimpanzee

Our next step was to perform a comparative quantitative analysis of the ChIP-seq read counts in the orthologous TEs that overlap H3K9me3 regions. By doing so, we aimed to comparatively characterize the degree of orthologous TE silencing in the two species. We classified TEs as putatively silenced only (or more strongly) in either humans or chimpanzees, when we found evidence for differences in H3K9me3 ChIP-seq read counts across the species (FDR < 1%; Supplemental Table 2). When we could not reject the null hypothesis that similar numbers of H3K9me3 ChIP-seq reads are mapped to an orthologous TE, we referred to the silencing status of these TEs as ‘shared’. That said, we recognize that inability to reject the null hypothesis does not provide strong evidence for the absence of a difference in silencing between the species.

As we reported above, we observed that TE overlap with H3K9me3 regions differs substantially across different TE classes (for example, 60% of SVA elements overlap H3K9me3 regions compared to only 10% of DNA transposons). Yet, the proportion of TEs for which we have evidence for inter-species differences in silencing is consistent, regardless of TE class (Fig. 4A). Indeed, while ultimately most TEs (88%) do not overlap H3K9me3 regions, when TEs do overlap H3K9me3 regions, we are generally (for 82% of TEs) unable to find evidence that these TEs are silenced differently across species. It is challenging to draw conclusions based on lack of evidence to reject a null hypothesis. Nevertheless, two lines of evidence support a conclusion that TE silencing status is generally conserved in humans and chimpanzees. First, we observed that different TE classes (with considerably different TE numbers) are generally likely to have similar shared or species-specific silencing status (Fig. 4B). Second, even when we use the adaptive shrinkage method (ashr; (Stephens 2017)), we find that at most, inter-species differences in TE silencing are associated with truly small effect sizes (with more than 80% of effect sizes smaller than 2 fold).

**Figure 4:**
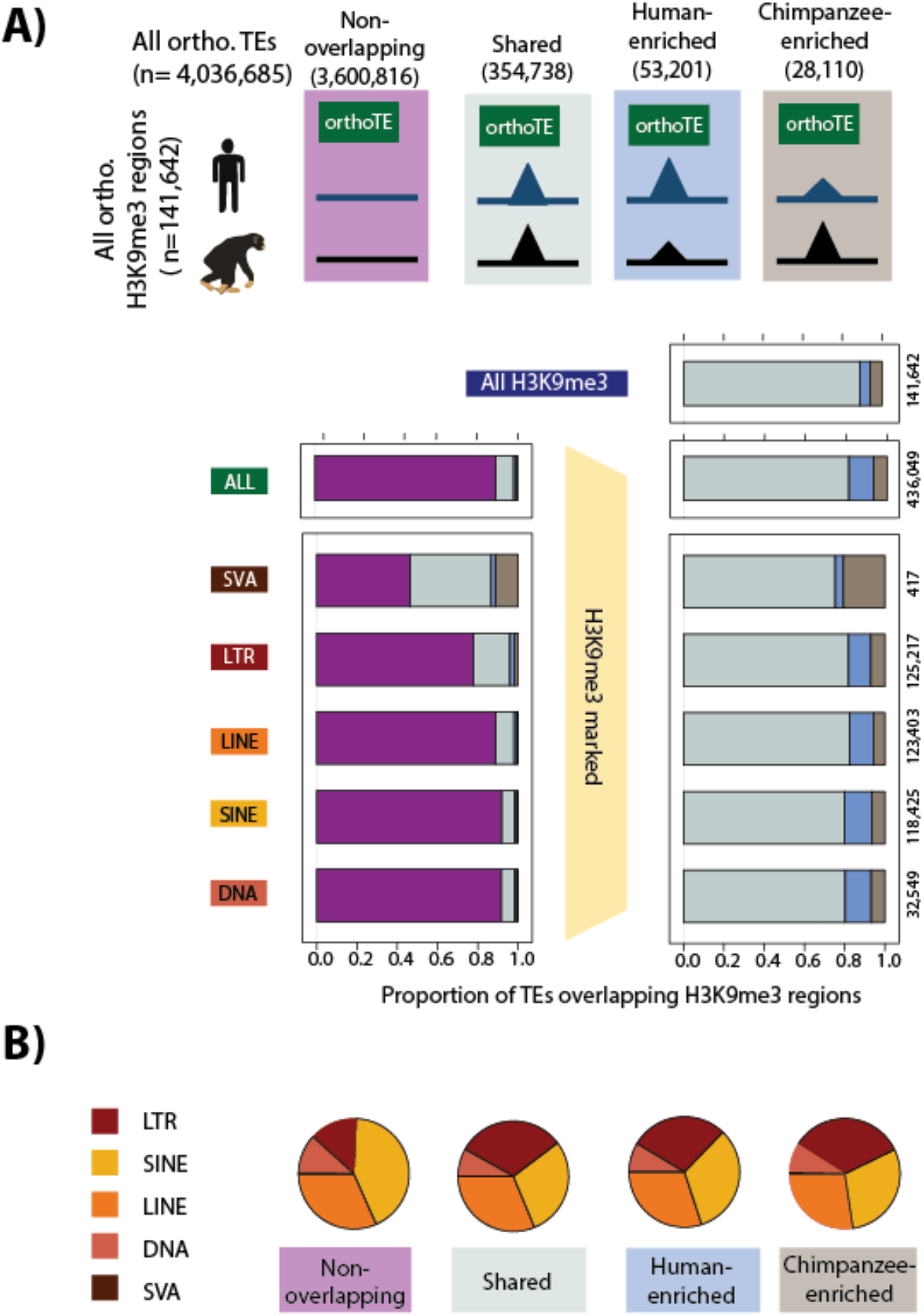
Majority of orthologous TEs may be similarly silenced in humans and chimpanzees. (A) The proportion of orthologous TEs in each class that do not overlap an orthologous H3K9me3 region (magenta), show similar H3K9me3 enrichment in both species (grey: Shared), are enriched in human (blue), or are enriched in chimpanzee (brown) are shown. Of those orthologous TEs that overlap an orthologous H3K9me3 region, the distribution between those that are Shared, Human-enriched, or Chimpanzee-enriched is shown in the right panel. (B) The proportion of orthologous TE elements belonging to each class within each of the four H3K9me3 silencing categories (LINE: orange, SINE: yellow, LTR: red, SVA: brown, DNA: peach). The small number of SVA elements means that this class is not visible in the pie chart.

We proceeded by considering a higher resolution classification of TEs to families, and then to types (Fig. 2). The observation that most TEs are silenced to a similar degree in both species is maintained at the family resolution as well (75-83%; Fig. S17). We then particularly focused on LTR and LINE TE types, given the previously reported association of these TEs with TRIM28- mediated silencing mechanisms (Castro-Diaz et al. 2014; Turelli et al. 2014). Human LTRs are comprised of HERV elements which can be alternatively classified into Class I (HERV-F,H,I,E,R,W families), Class II (HERV-K), and Class III (HERV-L) based on the original retrovirus that integrated into the genome (van der Kuyl 2012). Using a subset of these HERV types curated by Turelli et al., where the annotation is likely to be correct, we observed considerable heterogeneity in the frequency that different HERV families and types overlap H3K9me3 regions (Fig. S18A). However, we found that older Class III elements are less likely to overlap H3K9me3 regions than the more evolutionarily recent Class I and II elements (Pearson’s Chi-squared test; *P* < 0.001; Fig. S18B). This is consistent with the idea that older HERV elements no longer have active promoters, which need silencing.

In contrast, when turning our attention to mammalian-specific L1 LINE elements, we found that the youngest elements (L1PA2-L1PA8) overlap H3K9me3 regions less frequently than evolutionarily older elements (Pearson’s Chi-squared test; *P* < 0.001; Fig. S19A). This observation is consistent with a previous study, which reported that more evolutionarily recent L1 elements are bound by TRIM28 less frequently than older elements (Castro-Diaz et al. 2014). The unusual pattern whereby older L1 elements are more likely to overlap H3K9me3 is conserved in humans and chimpanzees (Fig. S19B).

### Silenced orthologous TEs have similar properties in human and chimpanzee

We next sought to identify properties that might distinguish TEs based on the following classifications: *(i)* TEs that do not overlap with an orthologous H3K9me3 region (non-overlapping); *(ii)* TEs that are associated with similar enrichment of H3K9me3 in both species (shared); *(iii)* TEs that are associated with significantly higher enrichment of H3K9me3 in humans *(iv)* TEs that are associated with significantly higher enrichment of H3K9me3 in chimpanzees.

We first considered the length of a TE, which may relate to its transposition competency and hence need for silencing. Indeed, on average, TEs silenced in at least one species are longer than those that do not overlap orthologous H3K9me3 regions (Wilcoxon-rank sum test; *P* < 0.001; Fig. 5). This observation is unlikely to be explained by chance alone given that our definition of overlap requires at least 50% of the TE length to overlap H3K9me3. Indeed, if we artificially increase the length of our non-overlapping TEs to match the median length of TEs marked by H3K9me3 in both species, we do not see any more overlap with H3K9me3 regions. We found the same pattern when we analyzed TEs by class, which in itself is associated with different lengths (*P* < 0.001; Fig. S20A). Given that each TE class represents a heterogeneous mix of TE families with different properties, we extended this analysis to the eleven most predominant TE families representing each of the five TE classes. Interestingly, across all TE families there is a strong correlation between the proportion of TEs that overlap H3K9me3 regions and the median length of the TEs in the family (Pearson’s correlation = 0.93; Fig. S20B-C). However, across TE families, we find a strong correlation between the median length of TEs that do not overlap H3K9me3 and those that do (Pearson correlation = 0.99; Fig. S20D). Similarly, we see a lower correlation when considering TEs of different length within a family (Pearson correlation < 0 .86; Fig. 20E). We therefore propose that longer TEs are silenced more often because they are more likely to have regulatory potential.

**Figure 5:**
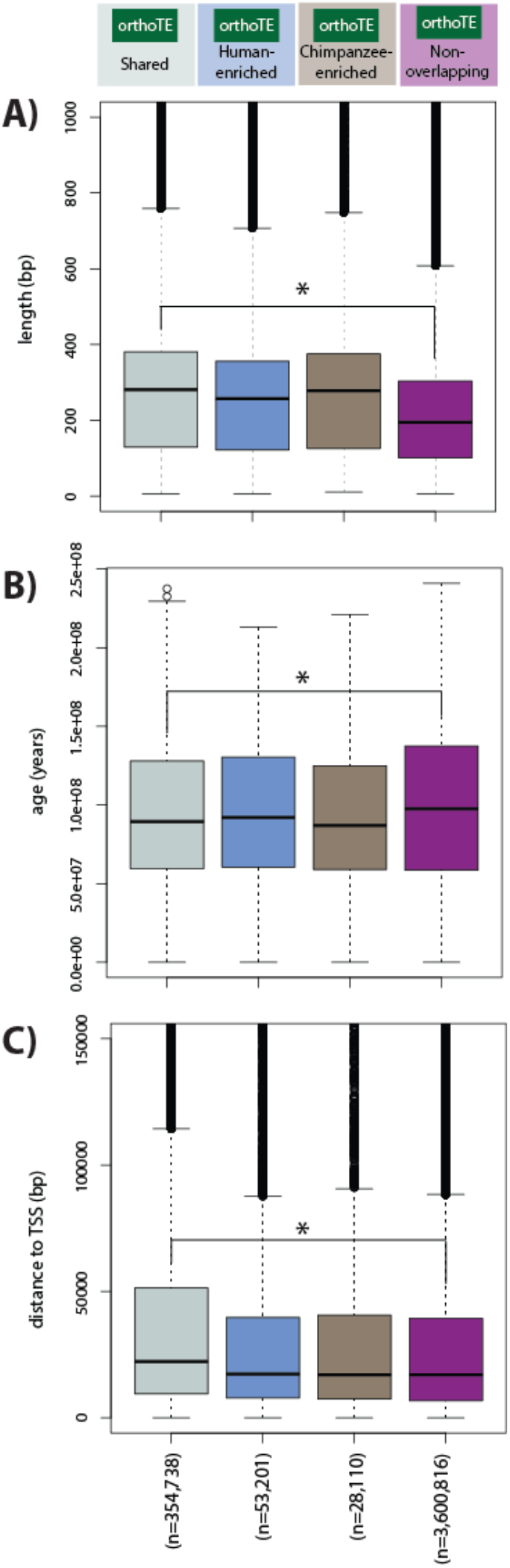
Majority of silenced orthologous TEs have similar properties in humans and chimpanzees. (A) The length of all orthologous TE instances occurring within each of the four H3K9me3 silencing categories. (B) The age of the TEs in the four categories. (C) The distance between each TE within the four categories and the closest orthologous TSS. Grey: Shared, blue: Human-enriched, brown: Chimpanzee-enriched, magenta: does not overlap an orthologous H3K9me3 region. Asterisk denotes when there is a significant difference between the Shared and Non-overlapping categories (Wilcoxon-rank sum test; *P* < 0.001).

We then asked whether TE copy number is correlated with silencing. Across TE families, we found a weak correlation between TE copy number and the probability of overlap with H3K9me3 regions (Pearson’s correlation = −0.47; permutation *P* = 0.15; Fig. S21). That said, in contrast to the general trend, TEs in the two smallest families (780 SVAs and 7,983 ERVKs) overlap H3K9me3 regions most often (Fig. S21).

We estimated the age of the TEs in each of the four silencing categories (see Methods). We found that TEs that are silenced similarly in both species are evolutionarily more recent (‘younger’) than TEs that do not overlap H3K9me3 regions (Wilcoxon-rank sum test; *P* < 0.001; Fig. 5). This observation is generally robust in most TE families, yet we found the opposite pattern when we focused on SVAs (Fig. S22). Across families, older TE families are associated with a lower proportion of TEs that overlap H3K9me3 regions (Pearson’s correlation = −0.81; Fig. S22).

Finally, we considered proximity of TEs to genes by determining the distance between each TE and the closest orthologous annotated Transcription Start Site (TSS). TEs that are silenced in both species are significantly further away from TSS than TEs that are preferentially silenced in one species, or those that do not overlap with H3K9me3 regions (Wilcoxon-rank sum test; *P* < 0.001 Fig. 5). This pattern is consistent across TE classes except for SVA elements, which are significantly closer to the TSS than other families (Fig. S23).

### Majority of orthologous TEs preferentially silenced in one species do not affect global gene expression divergence between species

To determine the potential impact of TE silencing on the regulation of neighboring genes, we characterized gene expression levels in the same iPSC lines from both species (using RNA-sequencing; see Methods, Fig. 6A, Fig. S24A, Table S2-3). After restricting the data to a set of human-chimpanzee-rhesus macaque orthologous exons (Blekhman et al. 2010) the RNA-seq data clusters primarily by species, and then by population (i.e. Caucasian or Yoruba / cell-type of origin; Fig. S24B). Indeed, after filtering lowly expressed genes, species emerges as the primary driver of variation in the RNA-seq data, as expected (Fig. S24C).

**Figure 6:**
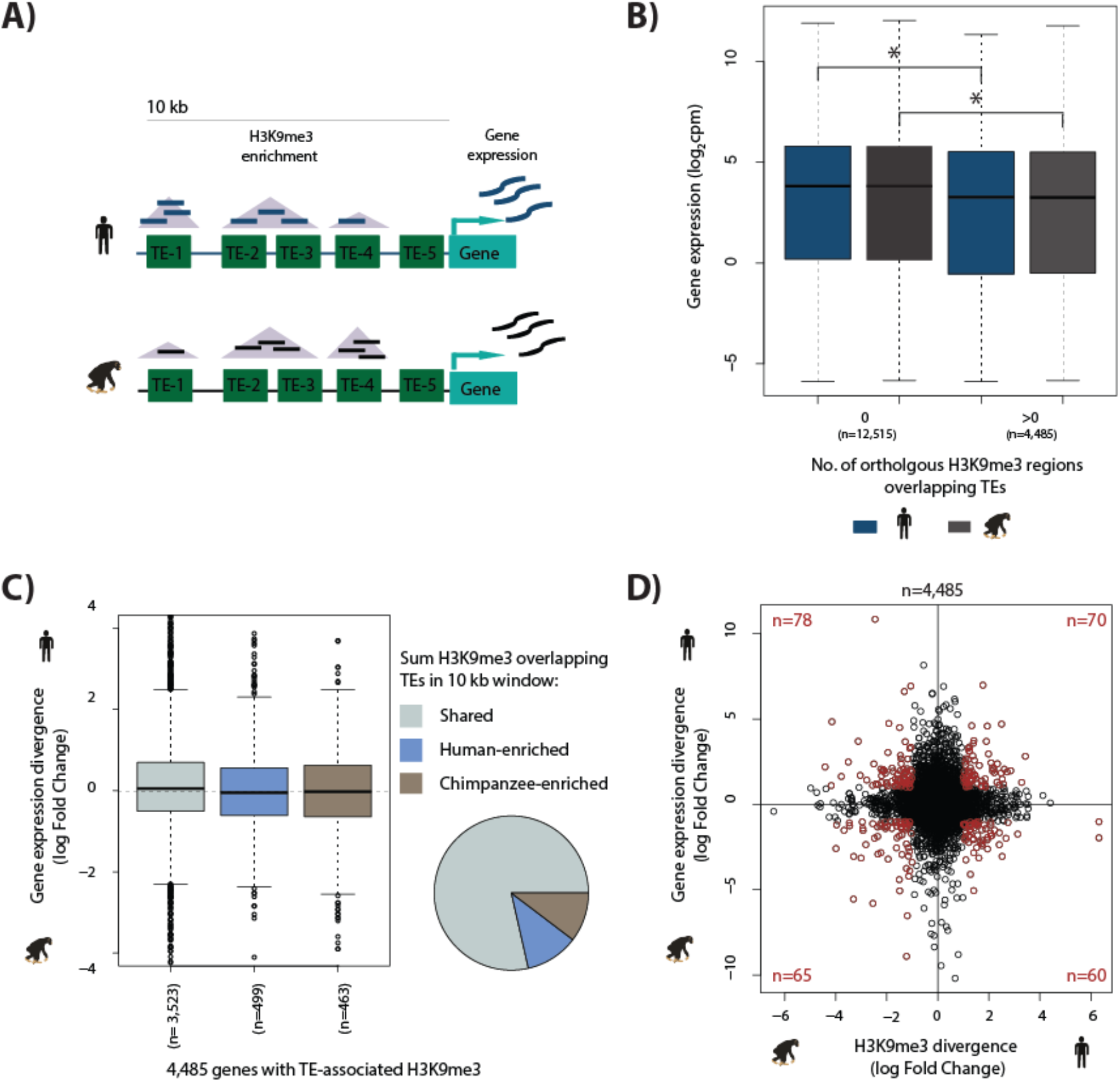
TE silencing divergence between species is not correlated with gene expression divergence. (A) Schematic representation of the experiment to determine how TE silencing affects nearby gene expression; orthologous TEs (green rectangles), orthologous H3K9me3 regions (purple triangles), human read counts (blue), and chimpanzee read counts (black). (B) Human and chimpanzee gene expression levels of orthologous genes with at least one TE present in a 10 kb window upstream of the TSS. Genes are categorized by the presence or absence of H3K9me3 regions overlapping TEs. (C) Gene expression divergence between human and chimpanzee at genes where TE silencing in the upstream 10 kb region is classified as Shared, Human-enriched or Chimpanzee-enriched. The proportion of genes associated with each TE silencing category is represented by the pie chart. (D) Divergence in H3K9me3-mediated TE silencing between species versus neighboring gene expression divergence. Genes which have a H3K9me3 log fold divergence of >1 or <-1, and a gene expression divergence log fold change of >1 or <-1 are shown in red. Asterisk denotes a significant difference between categories (Wilcoxon-rank sum test; *P* < 0.01).

To characterize the level of TE silencing associated with each orthologous gene’s expression, we defined a *cis*-regulatory region 10 kb upstream of the TSS of each orthologous gene that includes at least one annotated TE (Methods and Fig. 6A). As expected, we observed that gene expression levels decrease when TEs in the window overlap with H3K9me3 regions (Wilcoxon-rank sum test; *P* < 0.01; Fig. 6B). This observation is consistent with the general reported role of the H3K9me3 histone modification in gene regulation (Bilodeau et al. 2009; Mozzetta et al. 2015). The effect of TE silencing on gene expression is recapitulated when extending the window size to 20 kb, while TEs silenced within 1 kb of the TSS have a stronger effect on gene expression (Fig. S25). We then asked whether inter-species differences in H3K9me3 enrichment, specifically at TEs, are associated with gene expression divergence.

To obtain a quantitative measure of overall TE silencing in each species, we summed H3K9me3 ChIP-seq read counts from orthologous H3K9me3 regions overlapping TEs within 10 kb of annotated genes (see Methods). We used the same linear model framework described above to estimate inter-species difference in H3K9me3 counts, within a 10 kb window upstream of the TSS. Of the windows that contain at least one orthologous H3K9me3 region overlapping a TE, 79% have similar levels of H3K9me3 enrichment in human and chimpanzee (3,523), with 11% of regions enriched in human (499), and 10% of regions enriched in chimpanzee (463), at FDR of 1% (Fig. 6C). We then considered inter-species differences in expression levels for genes associated with similar or species-specific H3K9me3 enrichment at TEs.

We found no significant difference in the expression divergence effect size using 10, 20 or 40 kb windows upstream of the TSS (Fig. 6C and Fig. S26). A weak, negative association is observed when considering gene expression divergence of the 145 genes that show species-specific overlap of TEs with H3K9me3 within 1 kb of the TSS (Fig. S26). This effect on divergence at the TSS is similarly observed at the 539 orthologous genes whose TSS overlaps an H3K9me3 region irrespective of the presence of a TE (Fig. S27). Our observations are robust with respect to a wide range of FDR cutoffs used to classify inter-species differences in silencing, suggesting that incomplete power is not a likely explanation for the lack of association between divergence in silencing and expression levels (Table S6).

We performed an analogous analysis focused on inter-species divergence of H3K9me3 enrichment at TEs upstream of genes belonging to different categories. We similarly found that genes that are classified as differentially expressed between humans and chimpanzees, are not more likely to be associated with inter-species differences in H3K9me3 at TEs in their putative regulatory regions (Fig. S28A). Similar to the analysis described above, there is a relationship only when considering H3K9me3 overlapping TEs 1 kb upstream of the TSS. Again, the observation of no association of inter-species expression divergence and H3K9me3 enrichment in TEs is robust with respect to a wide range of statistical cutoffs used to classify either differential expression or differential ChIP-seq enrichment (Fig. S28B, Table S7). Indeed, when we considered the entire distribution, the correlation between inter-species effect sizes for gene expression divergence and H3K9me3 enrichment is weak (Pearson’s correlation = −0.0005 in a 10 kb window; the correlation in divergence increases to −0.14 when considering a 1 kb window upstream of the TSS; Fig. S26). Moreover, we observed nearly as many genes whose expression divergence is consistent with the observed patterns of TE silencing divergence (namely, TE silencing associated with lower expression level) as those that show the opposite, inconsistent pattern (Fig. 6D).

We next reasoned that perhaps TEs that are actively expressed in this cell type, are more likely to affect gene expression divergence. Indeed, TEs can be transcribed and contribute to lncRNA and neighboring gene expression (Karimi et al. 2011; Ramsay et al. 2017). We identified eleven TE types that are expressed in chimpanzee, and seven that are expressed in human (Methods; Table S8A). Notably, this analysis, in both species, identifies HERVH elements, which are known to be expressed in human pluripotent cells. Of the nine TE types that are expressed in at least one species, and annotated in both species, 1,025 instances are contained within our orthologous TE set (Table S8B). Only 123 of these instances overlap H3K9me3 regions, with 87 showing similar levels of H3K9me3 in both species, 14 enriched in human, and 22 enriched in chimpanzee (FDR of < 0.01). Only 19 genes showed species-specific silencing of a TE expressed in at least one species, within 10 kb of the TSS, we were thus unable to test the association between TE silencing/expression and gene expression divergence. In any case, the small number of instances suggests that gene regulation by an actively expressed TE is not a major mechanism of gene expression divergence.

We further hypothesized that perhaps imperfect annotation of orthologous TEs may contribute to the observation of lack of association between species-specific TE silencing and gene expression divergence. A clear association between silencing of non-orthologous TEs and gene expression divergence would provide some measure of support for this possible explanation. We thus performed a similar analysis focused only on the non-orthologous TEs. In this analysis we considered non-orthologous TEs that are located 10 kb upstream of annotated orthologous genes. We classified the non-orthologous TEs as ‘silenced’ or ‘not silenced’ based on all H3K9me3 regions identified in that species (we did not require H3K9me3 regions to be orthologous in this analysis). We found, once again, that there is no difference in gene expression divergence between genes with upstream non-orthologous TEs that are silenced, versus those genes with TEs that are not silenced (Fig. S29). A significant effect is only consistently observed when considering a 1 kb window immediately upstream of the TSS (Fig. S29). In other words, TE silencing, regardless of orthology, is not associated with gene expression divergence in human and chimpanzee iPSC lines.

As we mentioned we found our observations to be counter-intuitive. We therefore made one additional focused effort to identify species-specific regulatory effects of TE silencing by focusing on KRAB-ZNF genes. These genes, in addition to directing the repression of TEs, are themselves marked by H3K9me3 (O'Geen et al. 2007; Roadmap Epigenomics et al. 2015). We intersected a curated list of 256 KRAB-ZNF genes (Kapopoulou et al. 2016), filtered for orthology, with our list of orthologous H3K9me3 regions. We found that 104 (41%) of these genes overlap with orthologous H3K9me3 regions (Fig. S30A). The proportion of KRAB-ZNFs with similar H3K9me3 enrichment in both species (78% of overlapping genes) is equivalent to that observed for TEs (Fig. S30B). We found that the 256 KRAB-ZNF genes are no more likely to be differentially expressed at FDR < 1% between human and chimpanzee than all 17,354 genes (52% of KRAB-ZNFs are differentially expressed and 49% of all genes; Fig. S31A and Table S9). However, the 14 KRAB-ZNF genes whose TSS overlaps an H3K9me3 region that is enriched in either human or chimpanzee, irrespective of the presence of a TE, tend to associate with gene expression divergence between species (Fig. S31B).

## Discussion

The potential for TEs to participate in the regulation of gene expression has received increasing attention as genomic technologies to study these elements advance (Davidson and Britten 1979; Jordan et al. 2003; Feschotte 2008; Goke and Ng 2016). However, the role of TEs can be paradoxical: they can serve as regulatory sequence when they are bound by transcription factors, or localize in regions of open chromatin. Conversely, TEs can be actively silenced to prevent unwanted activity in a cell-type dependent manner. One cell type where TE activity can be particularly deleterious to genome integrity is embryonic stem cells. Indeed, it was found that TE activity is generally restricted through repressive histone modifications in mouse embryonic stem cells (Robbez-Masson and Rowe 2015; Schlesinger and Goff 2015). TE activity and regulation has been studied in many contexts, but there are no published comparative studies of TE silencing in primates.

In order to gain insight into how TE silencing evolves in pluripotent cells from the hominid lineage, we profiled the distribution of the repressive histone modification H3K9me3 in human and chimpanzee iPSCs. The rationale behind our approach is the expectation that TEs that are marked by H3K9me3 are silent and are therefore not mobile, nor able to act as regulatory elements in this cell type. In contrast, TEs that are unmarked and not silenced, could be expressed, or capable of transposition, or participation in gene regulation. Alternatively, these elements may have acquired sufficient mutations to lose the ability to transpose, or influence gene regulatory programs, and are therefore not silenced because they do not ‘threaten’ genome integrity. We were particularly interested in finding differences in silencing of the same TEs between species, as this could imply that these TEs act as regulatory elements in only one species, and could contribute to phenotypic differences between species.

Our balanced experimental design using episomally reprogrammed iPSCs from a panel of human and chimpanzee individuals, should have allowed us, in principle, to identify even modest differences in TE silencing between species. Our results, which we found a bit counter-intuitive, indicate that the degree and specificity of orthologous TE silencing in humans and chimpanzees are largely conserved. We acknowledge that proving the absence of a species-specific effect is challenging. In general, it is difficult to draw strong conclusions based on the inability to reject a null hypothesis. Nevertheless, our observations are robust with respect to a wide range of statistical thresholds used to identify TE silencing differences. There are other observations – patterns in the data - that support our conclusions, as we discuss below.

While TE silencing may be largely conserved, there are differences in the TE families that are preferentially silenced in both species. This suggests specificity of silencing and targeting mechanisms, which are shared in humans and chimpanzees. It also suggests that we have enough statistical power to detect these differences between TE families, which may indicate that we should have had sufficient power to detect inter-species differences as well. In our study, up to 60% of SVA, 50% of ERV, and 15% of L1 families overlap with H3K9me3 in both species. It is likely that TE silencing in our human and chimpanzee iPSC lines is mediated by the TRIM28 corepressor as it is known to be involved in silencing ERVs in mouse ESCs (Rowe et al. 2010), and SVAs, HERVs and L1 elements in human ESCs (Castro-Diaz et al. 2014; Turelli et al. 2014). This specificity is likely achieved through sequence-specific KRAB-ZNFs that recruit TRIM28. Indeed, SVA elements have been shown to be recognized by the KRAB-ZNF, ZNF91, and L1 elements by ZNF93 (Jacobs et al. 2014).

Perhaps surprisingly we did not see consistent trends with respect to silencing and the age of the TE. We also did not observe a clear relationship between the degrees or species-specificity of silencing and TE age (namely, TEs silenced preferentially in one species are not evolutionarily younger, as might be expected). Assuming an evolutionary arms race model, one expects that TEs and their KRAB-ZNF repressors would co-evolve; indeed there is a correlation between the age of KRAB-ZNF genes and their target TEs (Najafabadi et al. 2015), and related KRAB-ZNFs bind related TEs (Schmitges et al. 2016). However, the recent observation that many KRAB-ZNF gene – TE pairs arose at the same time during evolutionary history, and that this relationship is maintained to this day despite the fact that these TEs are no longer able to transpose, challenges the idea of a broad evolutionary arms race (Imbeault et al. 2017). In our study, we found silencing by H3K9me3 in TEs that are no longer transposition competent. It could be that these TEs are not silenced to prevent their mobility, but rather to prevent their potential regulatory activity.

TEs that are silenced in both species are located further away from the TSS than TEs that are silenced preferentially in one species. This difference between the ‘shared’ and ‘species-specific’ silenced TEs indicates that we do not have an overwhelming proportion of false negatively classified TEs in the ‘shared’ group. In other words, this is another observation supporting the significance of our inability to reject the null hypothesis in most tests. This observation also raises the possibility that TEs that are silenced only in one species are more likely to contribute to gene expression differences between species. However, we did not find this to be the case. Within species, genes with proximal TEs marked by H3K9me3 have lower expression levels, on average, than genes with proximal TEs that are not marked. Yet, across species, TEs silenced preferentially in one species do not have consistent effects on gene expression divergence between species. We were also not able to ascertain an effect on gene expression divergence when we restricted our analysis to non-orthologous TEs and their overlap with H3K9me3 in each species independently. We find these observations counter-intuitive and again – the basis for our inference is an inability to reject a null hypothesis, in this case, the inability to identify significant correlation between two data sets. Here, however, we were able to demonstrate quite clearly the robustness of our observations by performing multiple alternative analyses, as we report in detail in the results section.

We believe that our observations should be understood in the following context: (i) Even if our study is ultimately shown to be underpowered to detect very small differences in silencing and gene regulation, we can nevertheless conclude that the regulatory effects of TEs in our comparative system are not substantial. Other works agree: Even extreme perturbation by knockdown of the TRIM28 co-repressor in human neural progenitor cells, only leads to the up-regulation of 163 neighboring genes (Brattas et al. 2017). Overall, while studies have shown the effect of TE-derived enhancers on gene expression divergence at single loci (Sundaram et al. 2017), or tens of loci (Chuong et al. 2013), genome-wide effects on global gene expression divergence have been less clear. (ii) We have restricted our analysis to genes which are orthologous in sequence between the two species. It is possible that regulatory novelty co-occurs with gene novelty, though admittedly this is rare when the species considered are human and chimpanzee. (iii) It is likely that not all TEs have regulatory potential and hence need for silencing. Indeed, Imbeault et al. recently showed that KRAB-ZNF gene-bound TEs only affect neighboring gene expression if the TE overlaps marks of enhancers (i.e. H3K27ac and H3K4me1) in at least one cell-type. Overall gene expression levels were similar irrespective of whether the neighboring TE is marked by H3K9me3 (Imbeault et al. 2017). Further sub-setting TEs to those with potential regulatory activity could help identify those TEs that are likely to affect gene expression, and potentially gene expression divergence. Yet, arguably, this would not affect our general conclusions that overall, TEs contribute little to regulatory divergence in humans and chimpanzees. Our observations should also be interpreted in the context of our initial assumption about what can be learned from characterizing TE silencing. Indeed, the role, and contribution of TEs to gene regulation could analogously be assessed by characterizing TEs marked by active histone modifications (Lynch et al. 2011; Chuong et al. 2013; Ward et al. 2013; Xie et al. 2013; Villar et al. 2015; Trizzino et al. 2016; Sundaram et al. 2017), genomic location within accessible chromatin (Jacques et al. 2013), or expression as non-coding RNA (Karimi et al. 2011; Ramsay et al. 2017). In mammalian embryos, in particular, there is a complex relationship between TEs and the host genome where some elements need to be silenced, whilst the regulatory activity of other elements are integral to maintaining the pluripotent cell state (Schlesinger and Goff 2015; Gerdes et al. 2016; Izsvak et al. 2016). In our study we chose to focus on iPSCs as a model of the blastocyst inner cell mass, where TE activity is regulated by a histone-based silencing mechanism following the global resetting of DNA methylation. While there are examples of this TE repression pathway in differentiated cell types (Ecco et al. 2016), it is largely believed that DNA methylation replaces histone-mediated repression following the DNA de-methylation phase in embryogenesis. The ability of TEs to act as enhancer elements has been shown to be cell-type specific as characterized by cell-type specific chromatin enhancer marks and DNA methylation (Xie et al. 2013). Our results on repressive histone-mediated TE silencing may therefore not extend to other somatic cell types, which are not under the same level of regulatory constraint.

In summary, to date, there have been few genome-wide studies demonstrating the contribution of TEs to gene regulation in primates. TEs have been shown to comprise most primate-specific regulatory sequence (Jacques et al. 2013), to contribute to primate-conserved non-coding transcripts (Ramsay et al. 2017), and it has been suggested that exaptation of these elements contributes to lineage-specific gene regulation (Trizzino et al. 2016). Primates, however, may be an exception. Consistent with the reduction in TE activity in primates compared to other mammals, ChIP-seq assays for the highly conserved transcriptional insulator CTCF, have shown reduced TE-mediated propagation of CTCF binding sites in primates compared to other mammalian lineages (Schmidt et al. 2012; Schwalie et al. 2013). More broadly, while gene expression differences between primates were shown to often be associated with inter-species differences in histone modifications (Cain et al. 2011; Zhou et al. 2014), the chromatin landscape is generally highly conserved in primates. For example, as many as ~80% of genomic regions enriched for the active histone modification, H3K27ac, are similar between humans and chimpanzees in iPSC-derived neural crest cells (Prescott et al. 2015). Our results provide further support for the notion that chromatin state is generally highly conserved between closely related primate species. We have not found evidence that TEs, whether orthologous or not, are key drivers of gene regulatory evolution in humans and chimpanzees iPSCs.

## Methods

### iPSC culture

All chimpanzee and Caucasian (CAU) iPSCs were reprogrammed from fibroblasts with episomal vectors (Gallego Romero et al. 2015; Burrows et al. 2016), while the Yoruba (YRI) iPSCs were similarly reprogrammed from lymphoblastoid cell lines (LCLs) (Banovich et al. 2016). It has been shown, however, that cell-type of origin does not affect iPSC DNA methylation or gene expression patterns (Burrows et al. 2016). After reprogramming, iPSC colonies were cultured on a Mouse Embryonic Fibroblast feeder layer for 12-15 passages prior to conversion to feeder-free growth for 6-26 passages. Three new iPSC lines (H20682, H21792, H28815) were generated as previously described for H20961 (referred to as Ind1 F-iPSC in Burrows et al.), H28126 (Ind3 F-iPSC) and H21194 (Ind4 F-iPSC) (Burrows et al. 2016).

Human and chimpanzee feeder-independent stem cell lines were maintained at 70% confluence on Matrigel hESC-qualified Matrix (354277, Corning, Bedford, MA, USA) at a 1:00 dilution. Cells were cultured in Essential 8 Medium (A1517001, ThermoFisher Scientific, Waltham, MA, USA) at 37°C with 5% (vol/vol) CO_2_ with daily media changes. Cells were passaged by enzyme-free dissociation (0.5 mM EDTA, 300 mM NaCl in PBS), and seeded with ROCK inhibitor Y-27632 (ab120129, Abcam, Cambridge, MA, USA).

### iPSC quality control

All iPSCs were assessed for the expression of pluripotency markers, differentiation ability, and genomic instability: these cells express pluripotency factors, can spontaneously differentiate into all three germ layers following embryoid body formation, and display normal karyotypes. Gene expression from all samples was measured using the HumanHT12 Illumina Gene Expression Array, and data analyzed using the PluriTest bioinformatic assay to determine whether their global gene expression signature matches that of known pluripotent cells (Muller et al. 2011). We also assayed for the presence of episomal reprogramming vector sequence by qPCR. All three new iPSC lines (H20682, H21792, H28815), and previously described iPSC lines passed these quality control metrics (Gallego Romero et al. 2015; Banovich et al. 2016; Burrows et al. 2016). We identified one human (H28815) that tested positive for episomal reprogramming vector sequence. However, because this sample was not an obvious outlier in our data we chose to include it in our study.

### Immunocytochemistry

IPSCs were fixed in 4% paraformaldehyde in PBS, permeabilized in PBS-T (0.25% Triton X-100 in PBS), and blocked for 30 min in 2.5% BSA in PBS-T. Cells were incubated with primary antibodies in 2.5% BSA in PBS-T overnight at a 1:100 dilution: SSEA-4 mouse IgG (sc-21704, Santa Cruz Biotechnology, Santa Cruz, CA, USA), and Oct-3/4 rabbit IgG (sc-9081). Secondary antibodies were diluted to 1:500 in 2.5% BSA-PBS-T: donkey anti-mouse IgG, Alexa 488 (A21202, ThermoFisher Scientific), donkey anti-rabbit IgG Alexa 594 (A21207) and incubated for 1 hour at room temperature. Nuclei were counter-stained with 1 μg/ml Hoechst 33342 (H3570, ThermoFisher Scientific).

### ChIP-seq

ChIP-seq experiments were largely performed according to a published protocol (Schmidt et al. 2009). Briefly, 30 million cells were cross-linked with 1% formaldehyde for 10 min at room temperature followed by quenching with 2.5 M glycine. Following cell lysis and sonication (Covaris S2: 4 min, duty cycle 10%, 5 intensity, 200 cycles per burst in 4x 6 × 16 mm tubes per individual), lysates were incubated with 5 μg H3K9me3 antibody (8898, Abcam) overnight. ChIP and 50 ng input DNA from each individual was end-repaired, A-tailed and ligated to paired-end Illumina TruSeq ChIP Sample Preparation kit sequencing adapters before 18 cycles of PCR amplification. 200–300 bp DNA fragments were selected for sequencing. ChIP enrichment at known target genes (ZNF333 and ZNF554) was quantified by qPCR. ChIP-seq experiments were performed in two species-balanced batches. Input and ChIP libraries were multiplexed separately and 50 base pairs sequenced paired-end on the HiSeq2500 rapid run mode according to the manufacturer’s instructions.

Sequencing data quality was verified using FastQC (http://www.bioinformatics.babraham.ac.uk/projects/fastqc/). Paired-end ChIP-seq reads from each species were mapped to their respective genome (hg19 or panTro3) using BWA (version 0.7.12) (Li and Durbin 2009) and a mapping quality filter of MAQ >10. PCR duplicates were removed by samtools (version 0.1.19)(Li et al. 2009). Properly-paired reads only were selected for further analysis.

ChIP and Input reads from each individual were used to call peaks using MACS2 (Zhang et al. 2008) with the broadpeak option at various qvalue cut-offs. A lenient threshold of qvalue=0.1 was used in subsequent analysis to maximize the number of regions identified in each species.

To determine the minimum number of sequencing reads required to saturate the number of identified peaks, a read sub-sampling analysis was performed in two individuals from each species. This analysis revealed that the number of peaks called approaches saturation at a median read depth of 6.9 million paired ChIP reads across the four individuals tested (Fig. S7A). Subsampling input reads, while maintaining the number of ChIP reads constant, saturates the number of peaks called at a median of 5.7 million paired input reads across the four samples (Fig. S7B). 31/34 samples satisfied this minimum read number but as the three samples that did not meet this criterion represent both species we continued analysis with all individuals.

Peak co-ordinates in each species were then converted to the alternative genome using liftOver (Speir et al. 2016) and the best reciprocal chain (http://hgdownload.cse.ucsc.edu/goldenPath/hg19/vsPanTro3/reciprocalBest/) between hg19 and panTro3 requiring 70% sequence match. Peak regions that could be reciprocally and uniquely mapped between the two genomes were kept to generate a list of orthologous peak regions.

Orthologous peak regions in each species were concatenated to generate a final orthologous region set across both species. 50-mer mappability for every base in the hg19 and panTro3 genome was determined based on the GEM algorithm (Derrien et al. 2012). Mappability of all orthologous regions was determined, and only those regions with an average mapping quality score >0.8 in each species were maintained.

The number of reads falling into these orthologous peak regions was determined using HTSeq (Anders et al. 2015). Filtered, autosomal ChIP-seq read counts in each orthologous region (peak) from each individual in each species were compared using spearman correlation analysis. As sample H20961 was an outlier it was removed from further analysis. Counts were standardized and log transformed to generate log_2_cpm values for PCA analysis.

### Identifying differentially enriched H3K9me3-enriched regions

Orthologous regions with 0 counts in >8 individuals were removed. The R/Bioconductor package DESeq2 (Love et al. 2014) was used to identify regions differentially enriched between humans and chimpanzees. Regions with low mean read counts were filtered prior to multiple-testing correction. We controlled significance at FDR < 0.01 by the Benjamini-Hochberg method. Regions with a log_2_ fold change >0 and an adjusted p-value of <0.01 were considered to be ‘Human-enriched’, and those with log_2_ fold change <0 and an adjusted p-value of <0.01 as ‘Chimpanzee-enriched’. Those regions not significantly differentially enriched are defined as ‘Shared’.

Differential H3K9me3 enrichment across species was also assessed by transforming read counts by voom and identifying differentially enriched regions using limma (Law et al. 2014). Significance was controlled at FDR < 0.01.

The proportion of regions differentially enriched at a range of log2 fold changes was calculated both using DESeq2 output, and using an Empirical Bayes approach with adaptive shrinkage (ashr)(Stephens 2017).

### Orthologous TE generation

The RepeatMasker track (Smit, AFA, Hubley, R & Green, P. RepeatMasker Open-3.0.1996-2010 http://www.repeatmasker.org)(Jurka 2000) from the hg19 genome assembly (Speir et al. 2016) was converted to panTro3 genome coordinates using liftOver reciprocal chain files, requiring a 70% sequence match. These panTro3 coordinates were intersected with the panTro3 RepeatMasker track, requiring that 50% of base pairs overlap, and that the hg19-derived TE name matches that of the panTro3 RepeatMasker annotation. This set was subsequently lifted back over to hg19 again requiring a 70% match. The same procedure was followed starting with the panTro3 RepeatMasker track. Once RepeatMasker tracks were confidently obtained on the same genome, they were intersected (requiring 50% overlap). The distribution of orthologous TEs within the five main TE classes (LINE, SINE, LTR, DNA and SVA) is similar to the overall distribution of TE classes in each species (namely, when we do not require TEs to be orthologous; (Fig. S15)).

### Identifying silenced TEs

To obtain a high confidence set of TEs that are silenced, TEs that overlap H3K9me3-enriched regions by at least 50% of their length are considered to be ‘Overlapping’. We acknowledge that our stringent H3K9me3 region and orthologous TE definitions likely mean that young, active TEs will be excluded from the majority of our analyses.

### Non-orthologous TE analysis

The non-orthologous TE set for each species was generated after excluding the orthologous TEs from all RepeatMasker annotated TEs in that species. The list of species-specific TEs was generated by selecting non-orthologous TEs, where the TE name is not present in the RepeatMasker track of the other species. When determining the overlap between H3K9me3 regions and TEs with differing orthology status (orthologous, non-orthologous, and species-specific), we considered all H3K9me3 regions identified in each species independently (regardless of orthology).

### TE age determination

The age of each individual orthologous TE instance was estimated by dividing the number of substitutions from the consensus sequence (milliDiv score), obtained from the RepeatMasker track in human, by the human mutation rate (2.2 x10^-9^ per base per year)(Lander et al. 2001).

### Orthologous TSS annotation

A file containing annotated human transcription start sites (TSS) from hg19 was downloaded from UCSC (Karolchik et al. 2004). A single TSS was assigned per gene using the TSS of the 5’ most transcript from genes oriented on the sense strand, and the 3’ most transcript from genes on the anti-sense strand. Only TSSs corresponding to 30,030 genes contained in a primate orthologous exon file were retained (Blekhman et al. 2010).

### RNA-seq

RNA was extracted from ~3 million cells using the ZR-Duet DNA/RNA extraction kit (D7001, Zymo, Irvine, CA, USA) in two species-balanced batches. All RNA samples were of good quality: RIN scores of 10 were obtained for all samples except C40280 (9.7). All 17 samples were pooled together for RNA-seq library generation using the Illumina Truseq kit. The library pool was sequenced 50 base pairs, paired-end on the Hiseq2500.

Paired-end RNA-seq reads from each species were trimmed and mapped to each genome using Tophat2 (version 2.0.13) (Kim et al. 2013). Only properly-paired reads were taken for further analysis. The number of sequencing reads is similar across individuals from both species (median human: 42,557,599, median chimpanzee: 33,987,431)(Fig. S24 and Table S3). The number of reads falling into orthologous meta-exons across 30,030 Ensembl genes from hg19, panTro3 and rheMac3 (Blekhman et al. 2010) was determined using featureCounts within subread (version 1.5.0) (Liao et al. 2014). Autosomal genes with >0 counts in >10/17 individuals were retained. Counts were quantile normalized across all samples, standardized, and log-transformed to generate gene expression levels expressed as normalized log_2_cpm.

To identify genes differentially expressed between the two species, counts were transformed by voom (Law et al. 2014) prior to differential expression analysis using limma (Smyth 2004). We controlled significance at FDR < 0.01.

### Association between H3K9me3-mediated orthologous TE silencing and neighboring gene expression

We selected the annotated TSSs described above that correspond to orthologous genes for which we have gene expression data. We used the genomic coordinates of these annotations to define 1, 10, 20 and 40 kb windows upstream of the TSS. We then identified all TEs contained within each window, retaining the previous annotation of whether the TE overlaps an H3K9me3 region or not. Those TEs overlapping an orthologous H3K9me3 region had additional information in the form of H3K9me3 counts in each individual of each species. H3K9me3 counts in each orthologous H3K9me3 region overlapping a TE were summed within a window to yield a final H3K9me3 TE silencing count per gene per species. If multiple TEs fell within the same orthologous H3K9me3 region, this region was only counted once. H3K9me3 count values overlapping all TEs within a window was used as input for DESeq2 to determine inter-species differences in TE silencing per gene as previously described.

To determine gene expression divergence of genes overlapping H3K9me3 region at the TSS, regardless of the presence of a TE, a 1 kb region around the TSS was created. All 141,642 H3K9me3 orthologous regions (irrespective of the whether the regions overlaps a TE or not) were overlapped with the defined TSS corresponding to orthologous genes. Genes where at least 50% of the length of the 1 kb TSS region overlap with an H3K9me3 region are considered to be overlapping the TSS.

### TE expression

The expression of TE types was determined as previously described (Karimi et al. 2011). Briefly, multi-mapping, paired-end RNA-seq reads were retained for both humans and chimpanzees. RepeatMasker coordinates for all annotated instances of each TE type were obtained in each species. RPKM values for each TE type, in each species, were calculated by normalizing agglomerated read coverage of all annotated genomic copies of that TE type in the reference genome, to the agglomerated length of the TE type, as well as the total number of exonic reads. TE types were considered to be expressed if they had RPKM values >1.

### Association between H3K9me3-mediated non-orthologous TE silencing and neighboring gene expression

In each species independently, we used all previously determined non-orthologous TEs annotated as overlapping or not overlapping any H3K9me3 region identified in that species. Non-orthologous TEs were overlapped with the 1, 10 and 20 kb windows identified upstream of each orthologous gene’s TSS as described above. To identify TSS positions in the chimpanzee genome, the 1 bp TSS annotations of orthologous genes determined in human were lifted over to the chimpanzee genome as previously described in the generation of orthologous TEs. Similarly, a 1 kb region around each human TSS was lifted over to the chimpanzee genome, and only orthologous single bp TSSs where the broader 1 kb region lifts over to the chimpanzee genome were retained.

Each gene was categorized as ‘silent’ if all non-orthologous TEs in the window upstream of the TSS overlap H3K9me3, or ‘non-silent’ if at least one non-orthologous TE in the upstream window did not overlap with a H3K9me3 region. Results are consistent if we further stratify categories of genes where upstream TEs are fully silenced (all TEs overlap H3K9me3), or non-silenced (no TEs overlap H3K9me3).

### KRAB-ZNF gene overlap with H3K9me3 and expression

To identify those KRAB-ZNF genes marked by H3K9me3, a curated list of 346 genes H3K9me3 and expression (Kapopoulou et al. 2016) was filtered to include only those contained within our set of orthologous genes. These 256 genes were overlapped with orthologous H3K9me3 regions. KRAB-ZNF genes where 50% of their length overlaps with H3K9me3 are considered to be overlapping.

KRAB-ZNF genes where 50% of the length of a 1 kb region around the TSS overlaps with an orthologous H3K9me3 region were considered to be overlapping H3K9me3 at the TSS, and used in downstream gene expression divergence analysis.

## Supplemental Information

### Supplementary Figures and Tables

Figure S1-S31

Table S1-S9

### Supplemental Table 1

Table with 141,642 orthologous H3K9me3 regions, their ChIP-seq read counts, *DESeq2* results, and classifications as Shared, Human-enriched and Chimpanzee-enriched.

### Supplemental Table 2

Table with 4,242,188 orthologous TEs, their overlap with orthologous H3K9me3 regions, and classification as Shared, Human-enriched and Chimpanzee-enriched.

## Data Access

All data have been deposited in the Gene Expression Omnibus (www.ncbi.nlm.nih.gov/geo/) under accession number GSE96712.

## Acknowledgements

We thank Matthew Lorincz as well as all members of the Gilad lab, especially Sebastian Pott, for helpful and insightful discussions. We also thank Amy Mitrano for performing RNA extractions and preparing sequencing libraries. M.C.W is supported by an EMBO Long-Term Fellowship (ALTF 751-2014) and the European Commission Marie Curie Actions. This work was funded by an NIH grant from NIGMS (GM077959).

## Contributions

M.C.W conceived the study. M.C.W and Y.G designed the study. M.C.W performed experiments. M.C.W, S.Z, K.L and M.M.K analyzed data. B.J.P generated the iPSC lines. M.C.W and Y.G wrote the manuscript. M.S and Y.G oversaw the work.

